# Differential epigenetic landscapes and transcription factors explain X-linked gene behaviours during X-chromosome reactivation in the mouse inner cell mass

**DOI:** 10.1101/166249

**Authors:** Maud Borensztein, Ikuhiro Okamoto, Laurène Syx, Guillaume Guilbaud, Christel Picard, Katia Ancelin, Rafael Galupa, Patricia Diabangouaya, Nicolas Servant, Emmanuel Barillot, Azim Surani, Mitinori Saitou, Chong-Jian Chen, Konstantinos Anastassiadis, Edith Heard

## Abstract

X-chromosome inactivation (XCI) is established in two waves during mouse development. First, silencing of the paternal X chromosome (Xp) is triggered, with transcriptional repression of most genes and enrichment of epigenetic marks such as H3K27me3 being achieved in all cells by the early blastocyst stage. XCI is then reversed in the inner cell mass (ICM), followed by a second wave of maternal or paternal XCI, in the embryo-proper. Although the role of Xist RNA in triggering XCI is now clear, the mechanisms underlying Xp reactivation in the inner cell mass have remained enigmatic. Here we use *in vivo* single cell approaches (allele-specific RNAseq, nascent RNA FISH and immunofluorescence) and find that different genes show very different timing of reactivation. We observe that the genes reactivate at different stages and that initial enrichment in H3K27me3 anti-correlates with the speed of reactivation. To define whether this repressive histone mark is lost actively or passively, we investigate embryos mutant for the X-encoded H3K27me3 demethylase, UTX. Xp genes that normally reactivate slowly are retarded in their reactivation in *Utx* mutants, while those that reactive rapidly are unaffected. Therefore, efficient reprogramming of some X-linked genes in the inner cell mass is very rapid, indicating minimal epigenetic memory and potentially driven by transcription factors, whereas others may require active erasure of chromatin marks such as H3K27me3.

## Introduction

In mammals, dosage compensation between XX females and XY males is achieved by inactivating one of two X chromosomes during early female embryogenesis^1^. In the mouse, X-chromosome inactivation (XCI) occurs in two waves during early female development. The first wave takes place during pre-implantation development and is subject to genomic imprinting, resulting in preferential inactivation of the paternal X (Xp) chromosome^2^. In the trophectoderm (TE) and the primitive endoderm (PrE), which contribute respectively to the placenta and yolk sac, silencing of the Xp is maintained^3,4^. In contrast, in the epiblast precursor cells within the inner cell mass (ICM) of the blastocyst, (which correspond to mESCs), the Xp is reactivated and the second XCI wave and random inactivation of either Xp or the maternal X chromosome (Xm), occurs shortly after^5,6^. The inactive state is then stably maintained and transmitted through cell divisions in the soma.

Initiation of both imprinted and random XCI is dependent on the Xist long noncoding RNA (lncRNA) that coats the future inactive X (Xi) chromosome in *cis*. The essential role of *Xist* in initiation of imprinted XCI has been recently highlighted *in vivo* using single cell or single embryo allele-specific transcriptome analyses in early pre-implantation development^7,8^. Xist RNA coating is followed by gene silencing and several epigenetic changes, such as the depletion of active chromatin marks (*eg* tri-methylation of histone H3 Lysine 4 (H3K4me3), H3 and H4 acetylation), and recruitment of different epigenetic modifiers to the future Xi, including the polycomb repressive complex proteins PRC1 and PRC2, that result respectively in H2A ubiquitination and di-and tri-methylation of histone H3 Lysine 27 (H3K27me3)^9^. The inactive X chromosome is also enriched for mono-methylation of histone H4 lysine K20, di-methylation of histone H3 lysine K9 and the histone variant macroH2A^5,6,10^. Furthermore, our previous studies have shown that during the XCI process, different genes follow very different silencing kinetics^7,11^. Only during random XCI, in the epiblast, does DNA methylation of CpG islands occur to further lock in the silent state of X-linked genes, accounting for the highly stable inactive state of the Xi in the embryo-proper, unlike in the extra-embryonic tissues where the Xp is more labile^12–14^.

Much less is known about how the inactive state of the Xp is reversed in the inner cell mass (ICM) of the blastocyst. X-chromosome reactivation is associated with loss of Xist coating and repressive epigenetic marks, such as H3K27me3, from the inactive X^5,6^. Repression of *Xist* has been linked with pluripotency factors such as Nanog and Prdm14^15,16^. Studies on the reprogramming of somatic cells to induced pluripotency (iPSCs) have shown that X-chromosome reactivation required *Xist* repression and that occurs after pluripotency genes are expressed^17^. These observations suggest that the pluripotency program could enable X-chromosome reactivation via *Xist* repression as a first step. However, a previous study proposed that the reactivation of Xp-linked genes in the ICM operates independently of loss of Xist RNA and H3K27me3 based on fluorescent *in situ* hybridisation of nascent RNA (RNA FISH) and allele-specific RT-PCR analysis of a few (7) X-linked genes^18^. Therefore, it is still unclear whether X-chromosome reactivation in the ICM actually relies on pluripotency factors and/or on loss of epigenetic marks such as H3K27me3. Furthermore, whether loss of H3K27me3 is an active or a passive process has remained an open question. Given the speed of H3K27me3 loss on the Xp in the ICM from embryonic days 3.5 to 4.5 (E3.5-E4.5, ie 1-2 cell cycles), it is possible that active removal of the methylation mark may occur. Genome-wide removal of the tri-methylation of H3K27 may be catalysed by the JmjC-domain demethylase proteins: UTX (encoded by the X-linked gene *Kdm6a*), UTY (a Y-linked gene) and JMJD3 (encoded by *Kdm6b*)^19–22^. Diverse roles have been proposed for these demethylases^23–25^. JMJD3 appears to inhibit reprogramming^26^, whereas UTX plays a role in differentiation of the ectoderm and mesoderm^27^ and has been proposed to promote somatic and germ cell epigenetic reprogramming^24^. Interestingly, the *Utx* gene escapes from X-chromosome inactivation (*ie* is transcribed from both the active and inactive X chromosomes)^28^. This raises the intriguing possibility that Utx might have a female-specific role in reprogramming the Xi in the inner cell mass of the mouse blastocyst. *Utx* knockout mouse studies have suggested an important role of Utx during mouse embryogenesis and germline development, but its exact molecular functions in X-linked gene transcriptional dynamics have not been assessed^21,22,24,29,30^.

In this study we set out to obtain an in-depth view of the nature of the X-chromosome reactivation process in the ICM *in vivo*. We have defined the chromosome wide timing of X-linked gene reactivation and examined what the underlying mechanisms might be both at the transcription factor and chromatin levels. This work points to distinct mechanisms at play for the reactivation of X-linked genes in the ICM, with broad implications for our understanding of epigenetic reprogramming in general.

## Results

### Chromatin dynamics of the paternal X chromosome in the ICM of early to late pre-implantation embryos

Paternal X-chromosome reactivation has been described to occur in the pre-epiblast cells of pre-implantation embryos^5,6^, but the exact timing of X-chromosome reactivation and how the epigenetic marking of the inactive X chromosome (Xi) changes during this reprogramming is less clear. To determine the dynamics of chromatin changes on the Xp in the ICM during E3.5 (early) to E4.0 (mid) pre-implantation development, we performed immunosurgery on blastocysts at different stages in order to destroy outer TE cells and specifically recover the ICMs. By combining immunofluorescence with Xist RNA FISH, we analysed the enrichment of H3K27me3, as it is known to accumulate on the paternal inactive X shortly after the initiation of imprinted XCI, from E2.5 (16-cell stage)^5^. As previously reported^5,6^, H3K27m3 was found to be enriched on the Xist RNA coated X chromosome in almost all ICM cells of early pre-implantation blastocyst embryos (E3.5, 10-25 cells/ICM) (Figures 1a, b). Just half a day later, (E4.0, 20-40 cells/ICM), H3K27me3 enrichment and Xist RNA coating were lost from the Xp in approximately 25% of cells within the ICM (Figures 1a, b). The cells that lost Xist RNA coating and H3K27me3 enrichment on the Xp at E4.0 were often clustered together in close proximity in the ICM, suggesting that they represent the pre-epiblast population (Figure 1a). These results show that global Xp enrichment of H3K27me3 and Xist RNA coating are tightly correlated in the mouse blastocyst and that loss of this enrichment occurs with similar dynamics to loss of Xist coating in a subpopulation of ICM cells, presumably the pre-epiblast, between E3.5 and E4.0.

**Figure 1.**
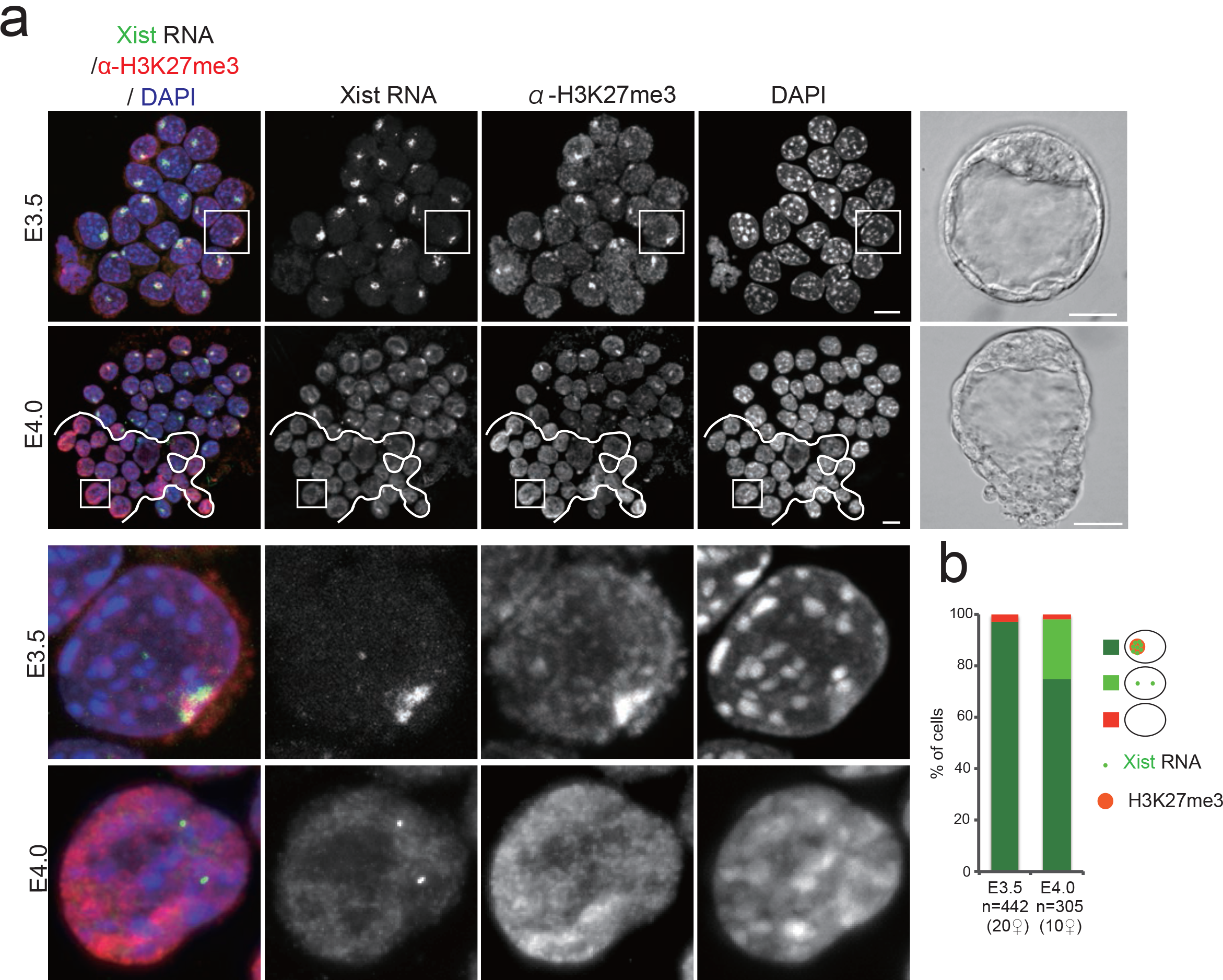
Xist RNA and H3K27me3 profiles in the ICM cells of early and mid blastocysts. **(a)** Examples of individual ICM of early (E3.5) and mid (E4.0) implantation stage embryos (photographs, scale bar 20µm) analysed by immunolabelling with antibodies against H3K27 tri-methylation (red) combined with Xist RNA FISH (green). For each stage, an intact ICM (IF/RNA FISH) and an enlarged nucleus are shown (scale bar, 10µm). The cells below the white line illustrate the cluster of cells that have lost Xist RNA coating and H3K27me3 enrichment on the Xp and are presumably the epiblast. **(b)** Proportion of ICM cells showing enrichment of H3K27me3 on the Xist RNA coated X chromosome in early and mid blastocyst stages are presented as mean. (right panel). Below the graph the total cell number analysed is indicated, followed by the total number of female embryos analysed in brackets. ICM, inner cell mass; RNA FISH, RNA-Fluorescent In Situ Hybridization; IF, Immuno Fluorescence.

### Different X-linked genes show different timing of reactivation and lineage-specificity

A previous report based on RNA-FISH and RT-PCR had shown that reactivation of seven Xp-linked genes seems to initiate despite the presence of Xist RNA coating and H3K27me3 enrichment of the Xp in ICM cells at E3.5 (early stage blastocysts)^18^. Combining RNA FISH and anti-H3K27me3 immunofluorescence, we analysed expression of two of these genes, *Rnf12* and *Abcb7* (both examined in the Williams et al., 2011 paper^18^) that are repressed during imprinted XCI by E2.5^7,11,31^. Strikingly, *Abcb7* and *Rnf12* showed very different reactivation behaviours in the early, E3.5, ICM (Figure 2a). While *Rnf12* exhibited low biallelic expression (<20% of ICM cells), suggesting its Xp silencing is maintained, *Abcb7* was biallelically expressed in almost all cells, despite the presence of Xist RNA coating and H3K27me3 enrichment on the Xp (Figures 2a, b, c). These results are only partially concordant with the Williams *et al.* study^18^, however the apparent discrepancy in *Rnf12* reactivation timing may be due to differences in the exact stages of blastocyst development examined, or to the different mouse strains used (B6D2F1 and B6xCast here, compared to CD-1 and CD-1xJF1 in Williams *et al*^18^).

**Figure 2.**
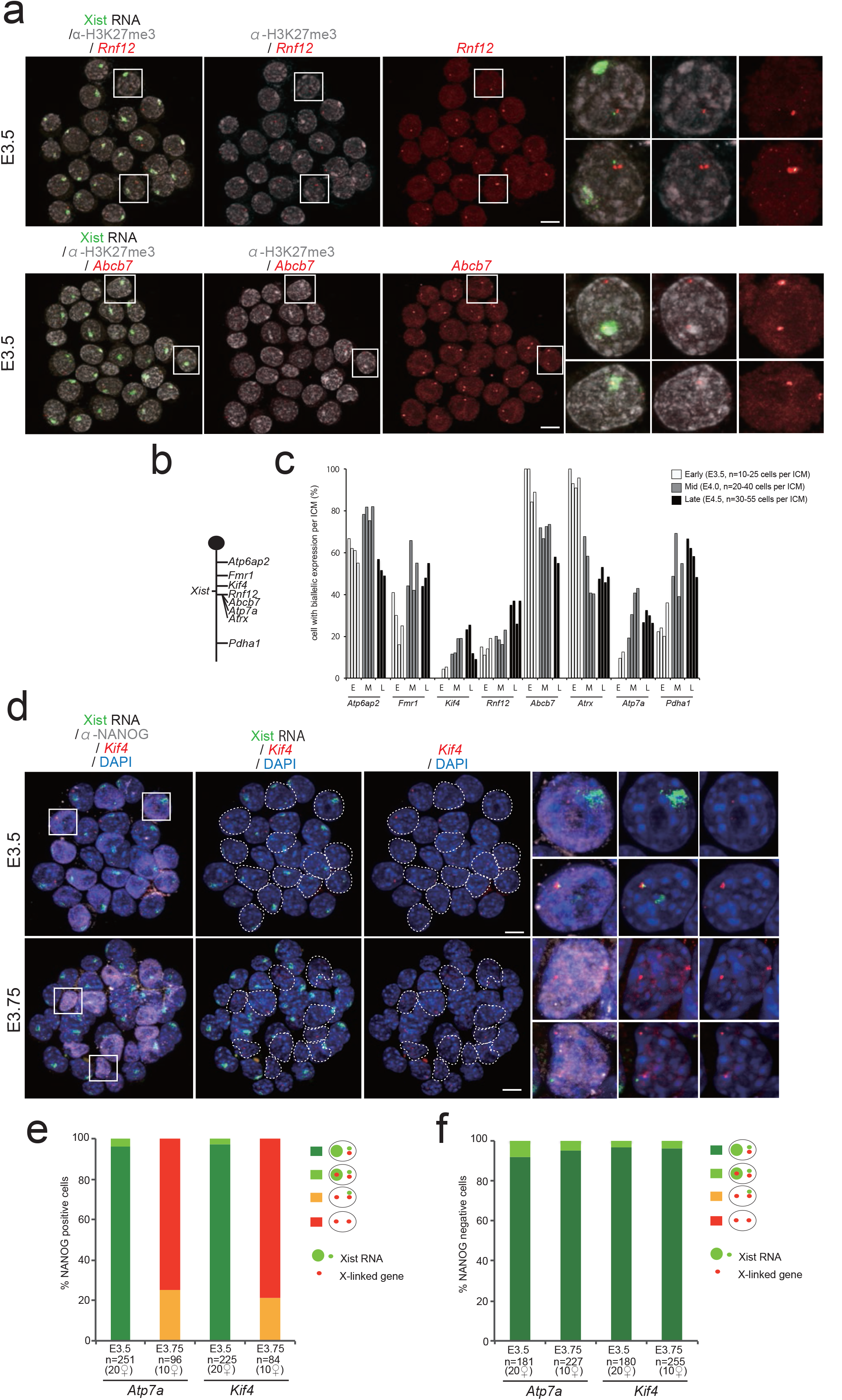
Xist RNA, X-linked gene expression and H3K27me3 profiles in the ICM cells of early to late blastocyst stage embryos. **(a)** Examples of individual ICM analysed by immunolabelling with antibodies against H3K27 tri-methylation (greyscale) and combined with RNA FISH for Xist RNA (green) and primary transcription from the X-linked genes (red), together with representative nucleus are shown (scale bar, 10µm). **(b)** Schematic representation of the X chromosome showing the location of the loci analysed in the panel **(a)** and **(c)**. *Atp6ap2* gene is known to escape XCI in 60% to 80% of blastocyst cells^11^ and used as a control of the experiment. **(c)** Percentage (mean) of cells showing biallelic expression for X-linked genes in ICM of independent early (E3.5), mid (E4.0) and late (E4.5) blastocyst stage embryos. **(d)** Examples of individual ICM analysed by immunolabelling with NANOG (greyscale), combined with RNA FISH for Xist (green) and X-linked genes (*Atp7a* and *Kif4*) (red) at early (E3.5) and mid (E3.75) blastocyst stage embryos. For each stage, an intact ICM (IF/RNA FISH) and enlarged nuclei (white squares) are shown. Dotted lines indicate the position of NANOG-positive cells (scale bar, 10µm). **(e)** Proportion (mean) of NANOG-positive ICM cells showing different Xist and X-linked gene expression patterns at early (E3.5) and mid (E3.75) blastocyst stage embryos. Below the graph the total cell number analysed is indicated, followed by the total number of female embryos analysed in brackets. **(f)**Proportion (mean) of NANOG-negative ICM cells showing different *Xist* and X-linked gene expression patterns at early (E3.5) and mid (E3.75) blastocyst stage embryos. Below the graph the total cell number analysed is indicated, followed by the total number of female embryos analysed in brackets.

We examined further genes for their timing of Xi reactivation in the ICM. We performed RNA FISH in pre-implantation (E3.5, early) through to peri-implantation (E4.5, late) blastocysts for 8 X-linked genes together with Xist (*Atp6ap2, Fmr1, Kif4, Rnf12, Abcb7, Atrx, Atp7a* and *Pdha1*) (Figure 2b). The genes were chosen based on their known range of silencing kinetics during imprinted XCI in pre-implantation embryos, including genes silenced early (prior to E3.0 such as *Kif4, Rnf12*, *Atp7a, Atrx* and *Abcb7*), late (after E3.0 e.g. *Pdha1, Fmr1*), or that escape XCI (*e.g. Atp6ap2*)^7,11^. Amongst the candidates, *Rnf12, Atp7a, Abcb7* and *Pdha1* were all previously described as being reactivated at the mid blastocyst stage (E4.0)^18^.

Increased frequencies of biallelic expression were observed for most genes in female ICM cells from the E4.0 stage onward (*Fmr1, Kif4, Atp7a* and *Pdha1 and Rnf12)*, indicating that they have reactivated in a subset of ICM cells (presumably pre-epiblast cells) (Figure 2c). However *Atrx* displayed biallelic expression as early as E3.5, similarly to *Abcb7* gene (also shown in Figure 2a). Thus, reactivation of *Atrx* and *Abcb7* occurs in the early ICM cells prior to any lineage segregation between epiblast (Epi) and primitive endoderm (PrE) cells^32–34^. Interestingly, just half a day later at E4.0, a decrease in biallelic expression of these two genes was seen in 30% to 60% of ICM cells (Figure 2c). Previous studies have shown that X-chromosome reactivation occurs in epiblast cells^5,6^, whereas PrE-derived tissues maintain an inactive Xp^4^. The decrease we observed in biallelic *Atrx* and *Abcb7* expression at E4.0 and E4.5 in ICM cells could indicate that these genes are silenced again, presumably in future primitive endoderm cells. In the case of *Atp6ap2*, which is a gene that normally escapes from XCI, as expected, it was found to be biallelically expressed in 60 to 80% of ICM cells at all stages^11^ (Figure 2c). Taken together, our data suggests that the reactivation of X-linked genes occurs with very different timing in ICM cells during early to late blastocyst stages. Furthermore, we find that a subset of genes may be reactivated early on, but then become rapidly silenced again in a sub-population of cells, presumably destined to become PrE.

To examine whether biallelic expression of more slowly reactivated genes correlates with pre-epiblast differentiation (and thus NANOG protein), we performed NANOG immunofluorescence combined with RNA FISH for Xist and two such X-linked genes (*Kif4* and *Atp7a*) in ICM cells of E3.5 (early) and E3.75 (mid) pre-implantation stage embryos (Figure 2d). As expected from our previous RNA FISH (Figure 2c), we found that cells mostly displayed monoallelic expression of X-linked genes (*Kif4*, and *Atp7a*) at E3.5. And this was the case in both NANOG positive and negative cell populations (Figure 2d, 2e, and 2f). *Kif4* and *Atp7a* then showed reactivation at E3.75 (Figure 2c) and the biallelic cells are almost all NANOG positive (Figure 2e). Moreover, biallelic expression of these X-linked genes was always observed in the absence of a Xist RNA cloud in NANOG-positive cells (Figure 2e, 2f). These results corroborate previous observations that *Atp7a* is reactivated only in cells expressing Nanog^18^. Our results suggest that both Nanog expression and loss of Xist RNA coating are linked to biallelic expression of late reactivated genes, but that Nanog expression alone is not sufficient. Taken together, our data point to reactivation in a lineage-specific manner beyond the mid ICM stage for genes that are late-reactivated. They also reveal a lineage-independent reactivation of the early-reactivated genes at E3.5 ICM.

### Single cell RNA sequencing of early and late pre-implantation female ICMs

The remarkable diversity in X-linked gene reactivation observed above (Figure 2) prompted us to explore the Xp reactivation process on a chromosome-wide scale. Furthermore, given the mixture of cells in the ICM, some of which are destined to become PrE, while others will become Epiblast, we were interested to know whether reactivation or silencing maintenance of Xp-linked genes correlated with PrE factor (*eg*. Gata4 or Gata6) and/or pluripotency factor expression (*eg*. Nanog, Oct4, Sox2) at the single-cell level^35^. We therefore performed single-cell RNAseq (scRNAseq) on ICMs of E3.5 and E4.0 pre-implantation female hybrid F1 embryos as well as used published trophectoderm (TE) cells where imprinted XCI is maintained^7^. The F1 hybrid blastocysts were derived from interspecific crosses between *Mus musculus domesticus* (C57Bl6/J) females and *Mus musculus Castaneus* (Cast) males. Single cells from individual ICMs were collected and polyadenylated RNA amplified from each cell according to the Tang *et al* protocol^36^ (n=17 cells from E3.5 ICM, and n=23 cells from E4.0 ICM and n=3 cells from E3.5 TE as control of XCI, Supplementary Table 1). We first assessed the extent to which transcriptomes of single cells from early (E3.5) and mid (E4.0) blastocysts were associated with each other, using principal component analyses (PCA, Figure 3a). We found that E3.5 ICM cells still showed substantial heterogeneity compared to E4.0 ICM single cells, which clustered into two distinct groups. Nevertheless, some signs that 2 sub-populations are emerging could be seen at E3.5 for some ICM cells. This revealed that developmental stage (E3.5 versus E4.0) does not seem to be the primary source of variability but that lineage specification between primitive endoderm and epiblast precursor cells could be important. We then also performed PCA analyses (Figure 3b) based on the expression levels of known pluripotency and differentiation factors, listed in Figure 3c. As expected from previous studies, E4.0 ICM cells fall into two clearly separated groups, either the PrE (expressing marker such as Gata4 and Gata6) or the Epi (expressing marker such as Nanog and Prdm14)^32,33^. No strong association was observed in E3.5 ICM cells with the exception of a few cells (n=3 potential pre-PrE and n=1 potential pre-Epi at E3.5, Figure 3b), supporting the idea that PrE and Epi lineages begin to be specified but are still not clearly established at the transcriptional level in E3.5 stage ICMs, as previously reported^34^. Next we performed a correlation analysis for single ICM cells and a few trophectoderm cells as a control, based on the expression status of pluripotency and differentiation factors (Figure 3c). As shown in Figure 3c, we classified cells according to their developmental stage and pluripotency/differentiation factor status: E3.5_TE (Trophectoderm of pre-implantation blastocysts), E3.5_ICM (Pre-lineage Inner cell mass of early pre-implantation blastocysts), E4.0_PrE (Primitive endoderm precursor cells of late pre-implantation blastocysts) and E4.0_Epi (Epiblast precursor cells of peri-implantation blastocysts). This clearly supports a shift from still rather heterogeneous transcriptomes in E3.5 ICM cells, into two well-defined subpopulations of pre-epiblast and primitive endoderm cells in E4.0 stage ICMs.

**Figure 3.**
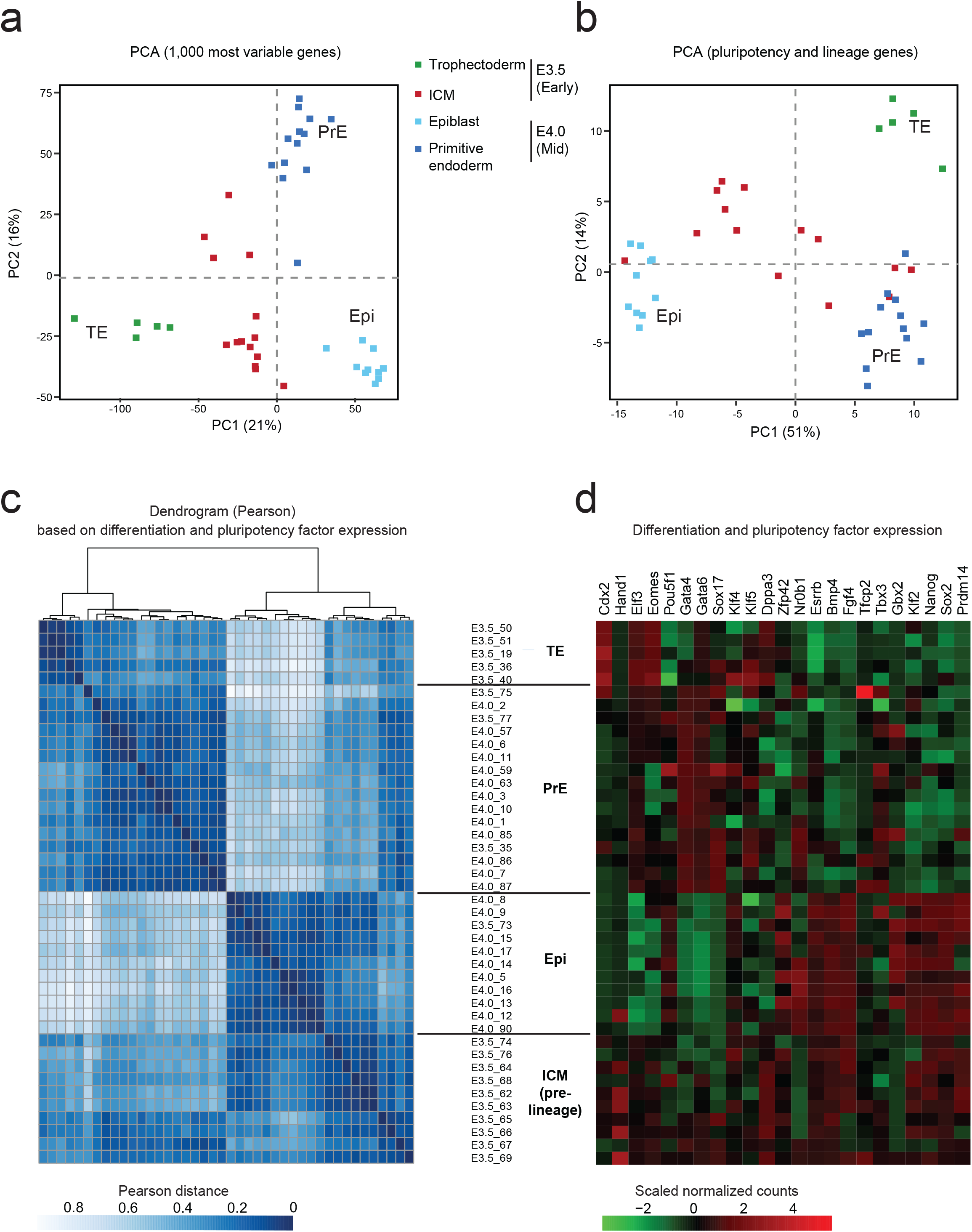
Single cell RNAseq reveals loss of heterogeneity in the E4.0 mid ICM compared to early E3.5 ICM. Principal component analysis (PCA) based on scRNAseq data from trophectoderm (E3.5), early (E3.5, 10-25 cells/ICM) and mid (E4.0, 20-40 cells/ICM) ICM cells on the 1,000 most variable genes **(a)** and on published pluripotency and differentiation candidate genes (n=23, list in Figure 3c) **(b)**. Different stages are designed by different colours. n= 14, 23 and 5 cells, respectively for E3.5 ICM, E4.0 ICM and E3.5 TE (details of each single cell is listed in Supplementary Table 1). **(c)**Hierarchical clustering (top) and Pearson distance (bottom) of pluripotency and lineage genes (listed in Figure 3d) expression variation in E3.5 and E4.0 single cells, based on Pearson’s correlation. Cells were clustered by lineage (TE, PrE and Epi), then by stage. n=42 single cell samples. TE, Trophectoderm; PrE, Primitive Endoderm; ICM, Inner cell mass; Epi, Epiblast. **(d)**Level of expression of the 23 candidate genes involved in pluripotency and lineage differentiation in the 42 single cell samples and used to classify cells according to their lineage are shown. Cells were ordered according to the hierarchical clustering in Figure 3c. TE, Trophectoderm; PrE, Primitive Endoderm; ICM, Inner cell mass; Epi, Epiblast.

### Specific X-linked gene behaviour highlighted by allele-specific analyses during X-chromosome reactivation

We next investigated chromosome-wide X-linked gene activity between early (E3.5) and mid (E4.0) ICMs. To assess the parental origin of transcripts, we took advantage of the high rate of polymorphisms between C57Bl6/J (maternal) and Cast (paternal) genomes that enabled us to distinguish Xm and Xp expression for informative transcripts (see Methods for expression thresholds and allele-specific pipeline). In this way an *in vivo* heatmap of X-linked gene activity was generated for early (E3.5_ICM) and mid (E4.0_PrE and E4.0_Epi) blastocyst stages and this was compared to trophectoderm cells at E3.5 (TE), extracted from the Borensztein et al. study^7^, as controls of X^P^CI maintenance (Figure 4a, Supplementary Figure 1 and Supplementary Table 2). To follow X^P^ reactivation by scRNAseq, we set a threshold of expression of RPRT=4 (Reads Per Retro-Transcribed length per million mapped reads, see methods) in at least 25% of the cells in both lineages (PrE and Epi) of mid blastocysts (n=116 genes), as in our previously published scRNAseq analysis of X-linked gene kinetics^7^. Low-expressed genes were excluded from the analysis in order to avoid amplification biases due to single cell PCR amplification. Trophectoderm cells (TE) from E3.5 female blastocysts were used as control for maintenance of imprinted XCI. They displayed 21% (18 out of 86) of biallelically expressed genes (allelic ratio >0.2, Figure 4a), and 17 of these genes are well-known escapees^7^. Interestingly, E3.5 ICM cells showed a higher number of biallelically-expressed genes when compared to TE. We found that 51% of X-linked genes were expressed from both X-chromosomes in E3.5 ICM (55 biallelic genes out of 107 in total, e.g. *Atrx*), despite the sustained expression of *Xist*. This supports our findings based on RNA FISH for early-reactivated genes (Figure 2) and further reveals the scale of such early reactivation. Intriguingly several of these reactivated genes (e.g *Atrx*, *Ubl4a* and *Eif1ax*) are clearly rapidly silenced again half a day later in PrE precursor cells only, as defined by the expression of 23 differentiation and pluripotency markers (e.g Gata4, Gata6 and Nanog, see Figure 3) (Figure 4a right panel). These data suggest that oscillations in the expression states of some genes on the Xp (such as *Atrx*) occur within a sub-population of ICM cells that will give rise to the PrE, where XpCI is known to be maintained, ultimately^4^. Our RNA-FISH data confirms that *Atrx* is transiently expressed from both X chromosomes even in the cells that will give rise to the PrE as it is found biallelically expressed in 90-100% of early ICM cells (Figure 2c). Our results reveal that there may be fluctuations in the inactive state of some genes during ICM progression, in the precursor cells of the PrE, rather than a straightforward maintenance of Xp silencing as previously thought.

**Figure 4.**
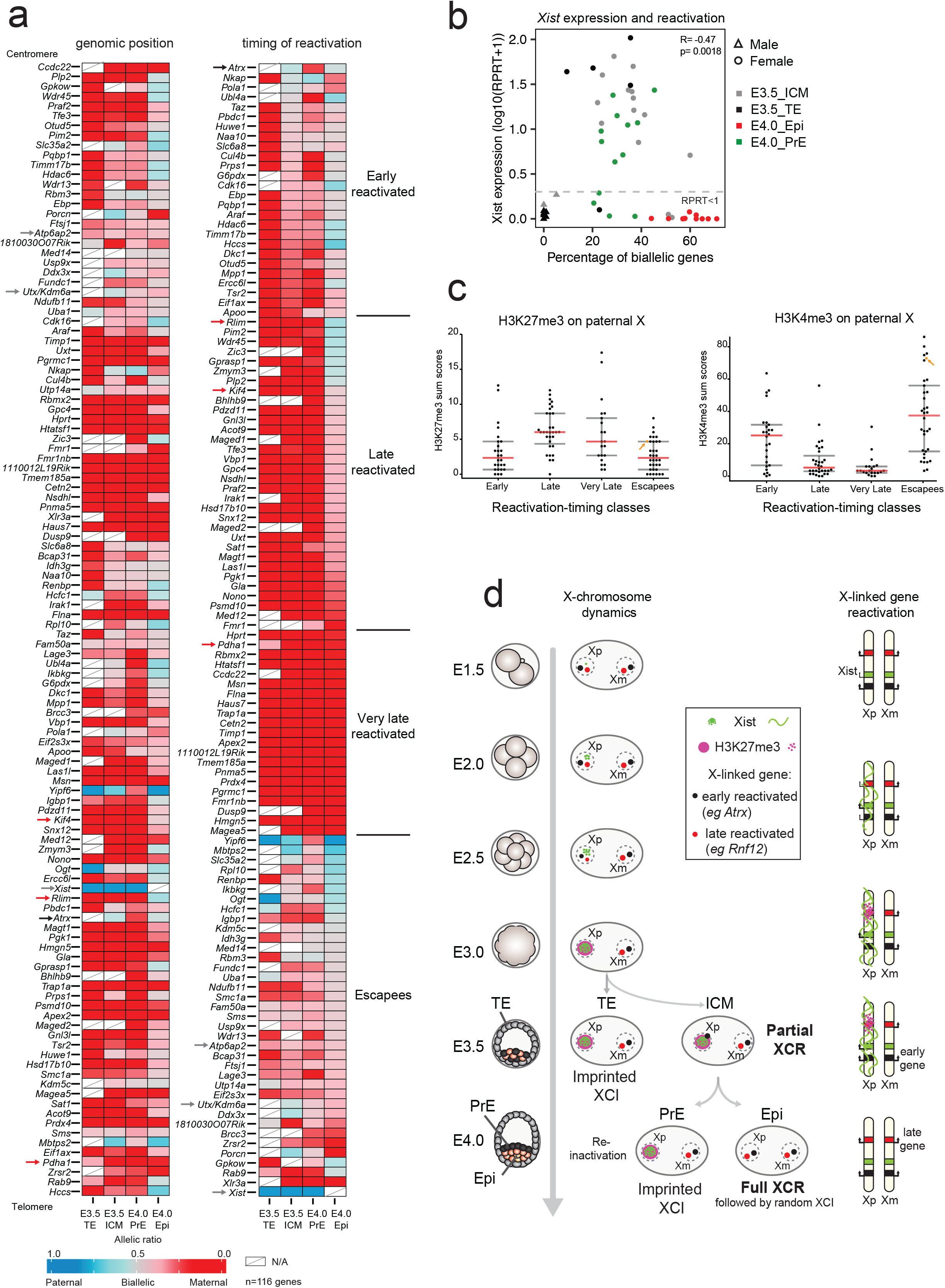
Different stages of X-linked gene reactivation in the ICM. **(a)**The mean of allele-specific expression ratios for each informative and expressed X-linked gene in E3.5 (Trophectoderm and ICM) and E4.0 (Primitive Endoderm and Epiblast) female B6xCast embryos are represented as heatmaps, with strictly maternal expression (ratio ≤0.15) in red and strictly paternal expression (ratio ≥0.85) in blue. Colour gradients are used in between these two values as shown in the key. Genes are ordered by genomic position (left) or by timing of reactivation (right). Further information is provided in Supplementary Table 2 and Methods. Blue, red and black arrows are respectively highlighting example of early, later reactivated genes and escapees. As expected, Xist RNA is paternally expressed in the trophectoderm cells. *Ogt* and *Yipf6* genes display similar paternal expression in the trophectoderm, escape imprinted XCI, and show random monoallelic expression and Castaneus bias respectively (Supplementary Figure 1)^7^. n= 116 genes. **(b)**Anti-correlation is shown between the level of Xist expression and the number of biallelically/reactivated and informative X-linked genes in scRNAseq (Spearman correlation). Male E3.5 single cells have been added and used as control for *Xist* expression and X-linked gene parental expression. Genes with level of expression as (RPRT<1) are considered as non-expressed in our samples. **(c)**Enrichment of H3K27me3 and H3K4me3 on paternal X chromosome obtained from (Zheng et al., 2016)^38^ shows significant differences (by Wilcoxon test) between Early and Escapee reactivation-timing classes compared to Late and Very Late. Low cell ChIPseq have been performed with ICM cells of pre-implantation embryos (pooled between E3.5-E4.0) after immunosurgery of the ICM^38^. Activated genes show an excess of H3K4me3 and repressed ones an enrichment of H3K27me3. *Xist* is highlighted with an orange arrow. Early versus Late (p=2.29*10^-4^ for H3K27me3 and p=1.63*10^-3^ for H3K4me3) and Very late (p=2.51*10^-2^ for H3K27me3 and p=3.95*10^-4^ for H3K4me3) and Escapees versus Late (p=1.95*10^-6^ for H3K27me3 and p=2.09*10^-7^ for H3K4me3) and Very late (p=7.33*10^-3^ for H3K27me3 and p=6.73*10^-8^ for H3K4me3). **(d)**Scheme of imprinted XCI, followed by reactivation in the inner cell mass of the blastocyst. Xp silencing is triggered by the long non-coding Xist RNA, followed by H3K27me3 recruitment. At the early blastocyst stage (E3.5), imprinted Xp is maintained in TE, when some genes are already showing reactivation in the ICM, independently of Xist (early reactivated genes). Those early genes are lowly enriched in H3K27me3 marks and highly enriched in H3K4me3 on their paternal allele compared to the later reactivated ones. Few hours later, when ICM cells are divided into PrE and Epi cells, Xp reactivation appears to be nearly complete only in the future Epiblast cells, accordingly to the loss of Xist and H3K27me3. In PrE, some early-reactivated genes could already be silenced again. This suggests a fluctuation of early-reactivated genes and different requirement of epigenetic memory between early and late-reactivated genes. TE, Trophectoderm; PrE, Primitive Endoderm; ICM, Inner cell mass; Epi, Epiblast.

In epiblast precursor cells, based on pluripotency factor expression (Figure 3), at E4.0, we noticed an absence of Xist expression and a marked progression in X_P_ chromosome reactivation as 77% of genes become biallelically expressed (89 out of 115, Figure 4a). Genes that showed reactivation only in epiblast precursor cells were classified as late-reactivated genes (Figure 4a) and confirmed our previous RNA-FISH data (*e.g. Rnf12*, *Kif4*, Figure 2). Interestingly, some genes classified as “very late reactivated” still appear to be repressed on the Xp, even at E4.0. In the case of *Pdha1*, this gene was found to be reactivated in about 40% of ICM cells at E4.0 by RNA-FISH (Figure 2c), compared to 4% of paternal expression in PrE and 18% in Epi, by scRNAseq (Supplementary Table 2). This could be explained by differences between nascent (RNA-FISH) and mature RNA (scRNAseq) for this gene, if the levels of paternal mRNA are not yet high enough for scRNAseq detection even though the gene has begun to be transcribed.

We describe here that X-chromosome reactivation can initiate for some genes independently of Xist loss and before lineage segregation at E3.5 (Figures 2a and 4a, Supplementary Table 2). However, in the epiblast precursor cells at E4.0, a higher percentage of biallelic X-linked genes was always observed in absence of Xist (Figure 4b). Indeed Xist expression levels and the percentage of biallelically expressed X-linked genes in single cells were anti-correlated (R=-0.47, p=0.0018, Spearman correlation). Thus, taken together, our data suggest that some genes (n=26 out of 116) undergo X-chromosome reactivation independently of Xist RNA and H3K27me3 loss, and that their expression fluctuates between early and mid ICMs, with many of them being re-silenced in the PrE lineage. The majority of X-linked genes will be reactivated later (E4.0), on the other hand, and only in epiblast precursor cells, in which pluripotency factors such as Nanog are expressed, and Xist RNA and H3K27me3 enrichment are lost (Figures 2 and 4).

### Differential TFs and H3K27me 3 enrichment in early and late reactivated genes

Next, we set out to define the features that are associated with the different categories of genes along the X as defined by their reactivation kinetics (early, late and very late or escapees). We assessed whether the timing of reactivation could be linked to the kinetics or efficiency of silencing of a particular gene. For this, we used our previously reported allele-specific scRNAseq analysis of imprinted XCI from the 2-cell stage to the early blastocyst (60-64-cell stage)^7^ and compared kinetics of silencing and timing of reactivation of X-linked genes. Correspondence analysis revealed that kinetics of reactivation of X-linked genes does not mirror their kinetics of silencing (Supplementary Figure 2a). Clearly the timing of reactivation is not simply about the lapse of time since silencing was initiated, nor about the location of a gene along the X chromosome (Supplementary Figure 2b). Although a slight tendency was observed for late and very late reactivated genes to be in close proximity of the *Xist* locus, *Atrx* and *Abcb7* genes are both silenced early, lie close to the *Xist* genomic locus and yet are also reactivated early^7,11,31^. Furthermore, our previous work revealed that although early silenced genes preferentially lie inside the first *Xist* “entry” sites as defined by Engreitz et al in ESCs, the late and very late reactivated genes failed to show any significant proportion correlation with *Xist* entry sites (Supplementary Figure 2c)^7,37^. Gene expression level was also not found as an obvious predictor of early or late reactivation (Supplementary Figure 2d). We thus hypothesize that late and very late reactivated genes may have acquired an epigenetic signature that prevents their rapid reactivation in early ICM cells, compared to early-reactivated genes. Early-reactivated genes on the other hand, may become expressed more rapidly due to specific TFs overriding their silent state.

We first examined recent allele-specific ChIPseq data for H3K27me3 and H3K4me3 in ICM of pre-implantation embryos (pooled between E3.5-E4.0)^38^. We overlapped the genes for which there is allelic information between this study and our different reactivation-timing groups and compared enrichment for H3K27me3 (left panel) and H3K4me3 (right panel) across their TSS (Figure 4c). We found a clear enrichment of H3K27me3 on the paternal allele but not on the maternal allele of late and very late reactivated genes compared to early-reactivated genes (respectively p=2.29*10^-4^ and p=2.51*10^-2^ by Wilcoxon test) and escapees (respectively p=1.95*10^-6^ and p=7.33*10^-3^ by Wilcoxon test) (Figure 4c left panel and Supplementary Figure 2e, left panel). Moreover, early-reactivated genes and escapees are enriched in the H3K4me3 histone mark compared to late (respectively p=1.62*10^-3^ and p=2.09*10^-7^) and very late genes (respectively p=3.95*10^-4^ and p=6.73*10^-8^) (Figure 4c and Supplementary Figure 2e right panel). As expected, we confirmed that H3K4me3-highly enriched genes are globally more highly expressed than lowly enriched genes (Supplementary Figure 2f). However as no association was found between a high level of could be a consequence of biallelic expression of early reactivated genes. Altogether, this highlights the asymmetric histone distribution between the different groups of genes during X-chromosome reactivation.

To explore the second hypothesis, that some TFs, including pluripotency factors, might drive expression from the Xp of a subset of early reactivated genes, we first analysed the correlation or anti-correlation between gene expression genome-wide and the degree of X-linked gene reactivation in female single cells (Supplementary Table 3, see Methods for details). As expected based on previous observations, Xp-chromosome reactivation correlates with pluripotency factors (e.g. Esrrb, Sox2, Nanog, Oct4 and Prdm14) and anti-correlates with PrE differentiation factors, such as Gata4, Sox17 and Gata6 ^5,6,15,16^. A gene ontology analysis of the top correlated genes (q-values <0.005) revealed that epigenetic modifiers are overrepresented (Supplementary Figure 2g) and corroborates our hypothesis that different epigenetic landscapes might at least partially underlie the different reactivation kinetics.

As X-chromosome reactivation is linked with epiblast formation and pluripotency gene expression, we then examined previously published datasets of transcription factor (TF) binding sites in mESCs (ChIPseq)^39,40^. In particular we analysed the occurrence of fixation sites at X-linked genes for pluripotency factors involved in Epiblast or embryonic stem cells differentiation (Nanog, Esrrb, Klf4, Oct4, Sox2, Tcfcp211 and Prdm14) and the Myc family also found associated with X-chromosome reactivation to a lesser degree (Supplementary Table 3). Indeed, Myc factors are expressed in early and mid ICM cells and there is a slight but significant association between high expression of *Myc* and *Mycl* genes and high rate of X-linked gene reactivation (Supplementary Figure 2h). Half of the X-linked genes, independently of their kinetics of reactivation and including escapees, presented at least one binding site for the above-mentioned pluripotency factors (data not shown). Their expression might be partially regulated by these factors^15,16^, but the binding of these factors alone cannot explain the behaviour of early reactivated genes. We next analysed for the presence of Myc family binding sites (Myc and Mycn binding sites, up to 3kb of the TSS and in gene body). Both escapees and early reactivated genes showed a surprisingly high enrichment for Myc factor binding sites, with respectively 42% (14 out of 33) and 31% (8 out of 26) showing at least one Myc binding site (Supplementary Figure 2i). In comparison, few late and very late reactivated genes displayed Myc binding sites with respectively 19% (6 out of 32) and 5% (1 out of 21) of them containing at least one binding site (p=0.0269 by Kruskal-Wallis). Myc transcription factors could thus be involved in transcription activation of silent X-linked genes and they have already been linked with the hypertranscription state described in ESCs and Epiblast^41^. Early reactivated genes and escapees could thus be targeted for reactivation from the silenced paternal X by the Myc TF family in early ICM.

In conclusion, the early reactivation of some X-linked genes, even prior to global loss of Xist RNA coating and H3K27me3 loss at E3.5, may be partly due to transcriptional activation thanks to Myc TF family, a lack of H3K27me3 and an enrichment of H3K4me3, while the majority of genes that are reactivated later show higher H3K27me3 and lower H3K4m3 enrichment, indicating a different epigenetic memory and response to TFs.

### Involvement of the histone demethylase UTX in efficient reprogramming of late reactivated genes

The above findings (Figures 2 and 4c) support a dependency between late and very-reactivated genes and loss of Xist and H3K27me3 from the Xp. To explore the hypothesis that epigenetic marking via H3K27me3 might play a role in the resistance of some genes to early Xp reactivation, we decided to impair H3K27me3 removal during the X-chromosome reactivation process. To do so, we produced peri-implantation (E4.5, late, n=30-55 cells per ICM) embryos lacking the X-linked histone demethylase UTX, which is reported to be specific for H3K27 demethylation^20–22^ and could promote reprogramming^24^. Interestingly *Utx* gene is expressed in pre-implantation embryos and remains high in early and mid ICM cells when it is down-regulated in trophectoderm (E3.5_TE), in which Xp inactivation is maintained (Supplementary Figure 3a). Homozygous knock-out mutant *Utx^FDC/FDC^* female embryos were obtained after matings between *Utx^FDC/Y^* studs (knock-out males) and *Utx^FD/+^*; or *Utx^FD/FD^*; *GDF-9iCre* females (Figure 5a) (FD = Flp and Dre recombined conditional allele; FDC = Flp, Dre and Cre recombined knockout allele (the GDF-9-driven Cre enables efficient recombination in the maternal germ line)^30^. Absence of UTX protein was validated by immunofluorescence at late blastocyst stage (E4.5) (Supplementary Figure 3b). Our aim was to assess if loss of Utx correlates with an accumulation of H3K27me3 at the inactive X, and its impact on the transcription status of two late-reactivated genes (*Kif4* or *Rnf12*). We performed immunostainings on E4.5 control, heterozygous and mutant female ICMs (Figures 5b and 5c). H3K27me3 enrichment on the Xp was retained in significantly more cells in *Utx* mutants compared to controls or heterozygous (respectively 73% versus 50% and 52%, p=0.0002, KW test). Furthermore a significantly higher proportion of Xist RNA negative cells with H3K27me3 enrichment was found in the mutant (5.1% vs 0.7% in controls, p=0.0067, KW test, Figure 5c and 5d). Altogether, our results are supportive of a scenario whereby UTX is actively involved in removal of H3K27me3 from the paternal X following Xist down regulation. Furthermore, when X-linked gene expression was assessed in the *Utx* mutant embryos, biallelic expression of *Kif4* or *Rnf12* were found with a corresponding absence of Xist, even in cells remaining H3K27me3 positive (Supplementary Figure 3c and 3d).

**Figure 5.**
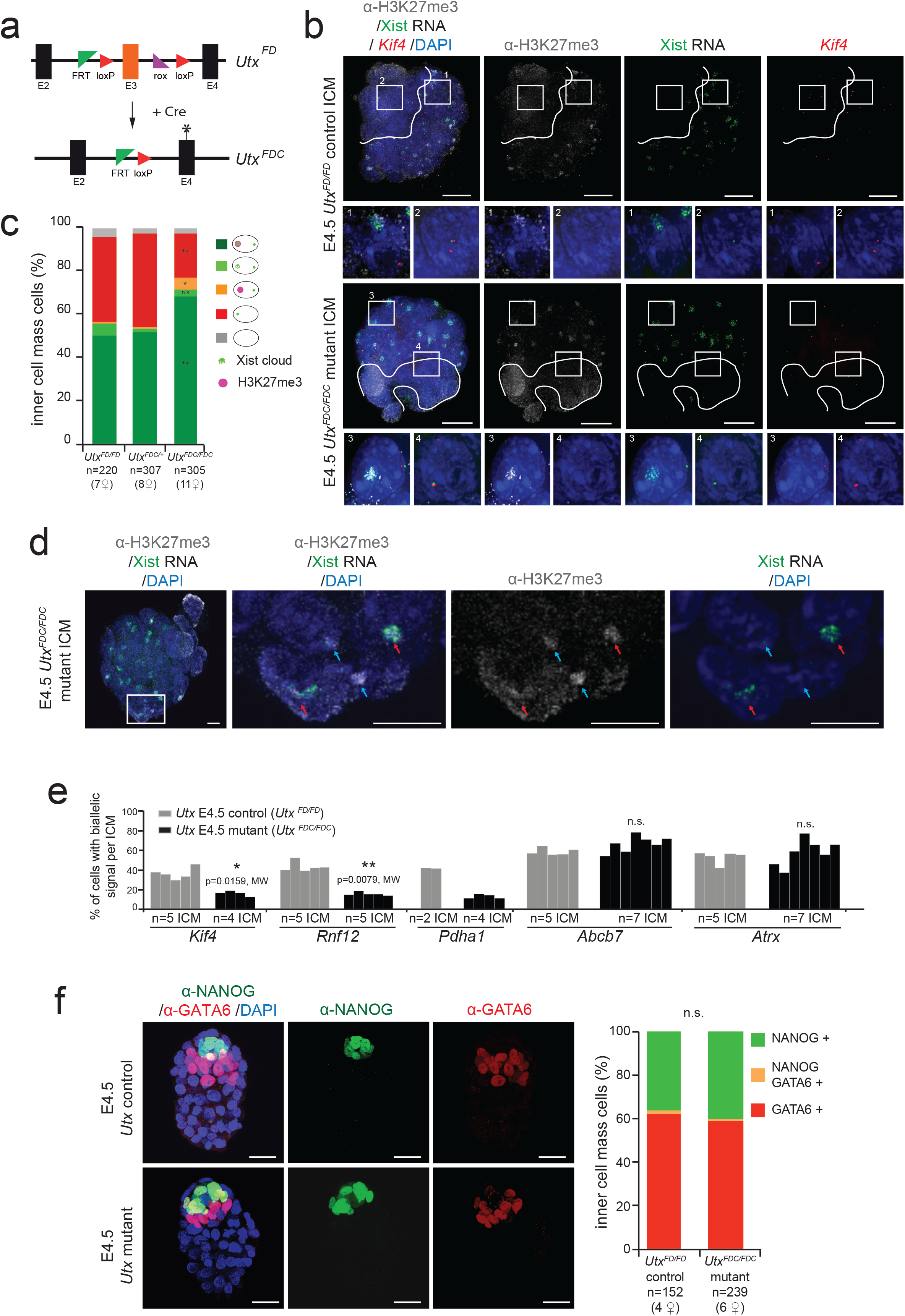
H3K27me3 UTX demethylase is required for proper reactivation of late-reactivated X-linked genes. **(a)**Conditional *Utx* allele: FD = Flp and Dre recombined conditional allele. Recombination of the third exon of *Utx* by Cre expression gives raise to a knockout FDC allele (the GDF-9 driven Cre enables efficient recombination in the maternal germ line). FDC = Flp, Dre and Cre recombined knockout allele. **(b)**Individual *Utx* control and mutant female ICM analysed by immunolabelling with H3K27me3 (grey), combined with Xist RNA (green) and *Kif4* gene (red) at late (E4.5) blastocyst stage. Enlarged nuclei are shown as example for not reactivated cells (1, 3) and reactivated cell (2, 4). The cells below the white line illustrate the cluster of cells that have lost Xist RNA coating and H3K27me3 enrichment on the Xp and are presumably the epiblast. Scale bars represent 20µm. **(c)**Proportion (mean) of ICM cells showing enrichment of H3K27me3 on the Xist RNA-coated X chromosome from E4.5 control *(Utx^FD/FD^*), heterozygous *(Utx^FDC/+^)* and mutant (*Utx^FDC/FDC^*) female blastocysts. Below the graph the total cell number analysed is indicated, followed by the total number of female embryos analysed in brackets. p-value<0.0057, <0.0018 and <0.021 between control and heterozygous versus mutant respectively for H3K27me3-Xist negative cells, H3K27me3-Xist positive cells and H3K27me3 positive, Xist negative cells, by two-sided Dunn’s test (Kruskal-Wallis and Post-hoc test). ** for p-value <0.01, * for p-value <0.05. **(d)**Second example of individual *Utx* mutant female ICM analysed by immunolabelling with H3K27me3 (grey), combined with Xist RNA (green) at late (E4.5) blastocyst stage. Red arrows pointed cells with both Xist and H3K27me3 enrichment on the Xp. Blue arrows pointed nuclei with only H3K27me3 enrichment on the Xp. Scale bars represent 10 µm. **(e)**Percentage (mean) of cells showing biallelic expression for X-linked genes in ICM of independent E4.5 control *(Utx^FD/FD^)* and *Utx* mutant (*Utx^FDC/FDC^*) embryos. *Kif4*, *Rnf12* and *Pdha1* are late or very-late reactivated genes, when *Abcb7* and *Atrx* are early-reactivated genes based on both IF/RNA-FISH and scRNAseq. MW, Mann-Whitney nonparametric test. **(f)**Maximum intensity projection of control *(Utx^FD/FD^)* and *Utx* mutant (*Utx^FDC/FDC^*) E4.5 blastocysts analysed by immunofluorescence against NANOG (green) and GATA6 (red). DAPI is in dark blue. Scale bars represent 20µm. Percentage of positive cells for Nanog, Gata6 or both have been assessed and summarized as the mean. Below the graph the total cell number analysed is indicated, followed by the total number of female embryos analysed in brackets. Non significant (n.s.) by Kruskal-Wallis test.

To explore the impact on gene reactivation further, we performed RNA-FISH on several late-or very-late reactivated genes (*Kif4*, *Rnf12* and *Pdha1*), as well as on early-reactivated genes (*Abcb7* and *Atrx*). Strikingly, *Rnf12*, *Kif4* and *Pdha1* reactivation was always lower in the mutant E4.5 ICMs compared to controls (about 50% decrease in *Utx* mutants, Figure 5e). Furthermore, this decrease correlated well with the increase in H3K27me3 positive cells in mutants. On the other hand, *Abcb7* and *Atrx* gene reactivation rates did not appear to be affected in *Utx* knockout embryos. Thus early reactivated genes do not appear to be sensitive to the lack of UTX and increase of H3K27me3 in ICM cells, supporting their H3K27me3-independent reactivation mechanism, as suggested by their depletion in H3K27me3 (Figure 4c).

Finally, to exclude the possibility that the apparent interference with X-chromosome reactivation in the pre-epiblast cells might actually be due to delayed or abnormal development in *Utx* mutant embryos, leading to an increased proportion of pre-primitive endoderm cells, we stained for NANOG (Epi marker) and GATA6 (PrE marker) in both control and mutant E4.5 female ICMs (Figure 5f). No difference was seen in the total number of cells per ICM (with a mean of 38, 40 and 40 cells per ICM respectively for *Utx^FD/FD^*, *Utx^FDC/+^* and *Utx^FDC/FDC^*) and in the proportions of NANOG (epiblast) and GATA6-positive (primitive endoderm) cells. Thus, the absence of Utx and subsequent retention of H3K27 methylation did not impact on ICM progression but does impair the efficiency of X-linked gene reactivation *in vivo*, at least for later reactivated genes. In summary, our results reveal the existence of different epigenetic memory states during imprinted X^p^CI, with some genes being sensitive to the requirement for Utx for removal of H3K27me3 and reactivation, whereas others can be reactivated independently of global Xist and H3K27me3 enrichment.

## Discussion

Transcriptional reactivation of the paternal X-chromosome occurs in the mouse ICM during pre-to peri-implantation development. The extent and nature of this reprogramming process has remained poorly defined until now. Our single cell analysis of paternal X-chromosome reactivation in the ICM provides the first chromosome-wide map of X-linked gene activity and strong evidence for multiple mechanisms involved in the loss of silencing of X-linked genes. Emergence of ICM, at the blastocyst stage, is a key event during early mouse development. We now know that pluripotency factors such as Nanog will be retained in the epiblast precursor cells that will give rise to the embryo-proper and this is where Xp-reactivation occurs^5,6,33^. At the early blastocyst stage (E3.5), primitive endoderm and epiblast precursor cells only begin to segregate and heterogeneity in the expression of specific lineage markers is still seen (*e.g*. Nanog and Gata6), as confirmed in our study (Figure 3). This initially high degree of cell-to-cell variation in pluripotency and lineage factor gene expression (*eg* Nanog, Gata6) is lost by E4.0, when two transcriptionally distinct populations of cells can be observed. The pre-Epi cells are characterised by pluripotency genes and loss of *Xist* expression; PrE cells show Xist expression, decreased pluripotency gene expression and enhanced lineage markers such as Gata4 and Gata6.

Early work showed that imprinted X-chromosome inactivation remains in extraembryonic tissues, including the yolk sac derived from primitive endoderm cells^3,4^ when Xp is reactivated in the pre-epiblast cells^5,6^. Previous studies have shown that loss of Xist RNA coating and H3K27me3 enrichment during X-chromosome reactivation was linked to pluripotency factors, such as Nanog and Prdm14^15,16^. The data we present here suggests that X-chromosome reactivation correlates with epiblast differentiation however reactivation of some genes is not limited to the future epiblast cells but initiates independently of lineage segregation in early pre-implantation blastocysts. Indeed, our IF/RNA FISH and scRNAseq analysis at E3.5 ICMs suggests that X-linked gene reactivation can initiate before loss of Xist and H3K27me3 and before the strict emergence of PrE and Epi precursor cells (Figures 2 and 4). This suggests that Xp chromosome reactivation and the pluripotency program can be uncoupled for some genes such as *Atrx* that are reactivated early on in almost all the cells of the E3.5 ICM. Importantly some of these early-reactivated genes then show Xp silencing again in E4.0 PrE. This implies a fluctuation in Xi status between E3.5 and E4.0, rather than a constant maintenance of Xp silencing, in future primitive endoderm cells (Figure 4d). Overall, our study highlights the distinct types of behaviour for different X-linked genes when it comes to X-chromosome reactivation. In the case of late-reactivated genes, reactivation is lineage-specific and restrained to the pre-Epi cells of the mid blastocyst onwards. Later gene reactivation shows a strong correlation with the presence of NANOG protein (Figure 2d) and with loss of Xist expression (Figure 4b) and H3K27me3 enrichment (Figure 2a). Moreover, loss of Xist RNA coating is the most predictive factor for biallelic expression of the late reactivated genes (Supplementary Figures 3c, 3d).

Our discovery that there are at least two different categories of X-linked genes in terms of their Xp reactivation behaviour is an important step in better understanding X-chromosome reactivation and epigenetic reprogramming in general. Interestingly, level of expression and genomic localization of X-linked genes are not obvious predictors of their reactivation behaviour (Supplementary Figure 2b and 2d). Our correlative analyses suggest that the dynamic presence of the Myc family of TFs might play a role in facilitating some early-reactivated genes to become re-expressed in ICM cells, but then revert to a silenced state in PrE cells (Supplementary Figure 1d and 1e). In the search for other TFs potentially involved in early X-linked gene reactivation, we used algorithms for motif discovery (see Methods). No specific TF binding motif associated with escapees and early reactivated genes could be found with a high confidence. The lack of enrichment for known motifs could be due to the limited number of genes included in each of the reactivation classes. However, motif comparison analysis of any over-represented motifs in escapees and early-reactivated genes revealed a correspondence with the transcription factor YY1 (Ying Yang 1), (p-value=0.0002). This motif occurs 2.5 times more frequently in the group of escapees and early reactivated genes (n=20, 57 promoters), than in the group of late and very late genes (n=7, 49 promoters). YY1 is associated with escapees in human and has previously been described to be co-bound to the same binding sites as MYC in mouse ESCs ^42,43^. The precise roles of the MYC proteins and YY1 in relation to Xp gene activity merits future exploration.

To better understand the degree to which epigenetic chromatin states might be involved in maintaining inactivity, we studied allele-specific H3K27me3 and H3K4me3 enrichment (Figure 4c and Supplementary Figure 2e). Distinct patterns of differential enrichment of these histone marks was found for early-reactivated genes and escapees (high K3K4me3 on the Xp) and later-reactivated genes (high H3K27me3 on the Xp). These different epigenetic signatures might underlie the distinct transcriptional behaviours of those genes during Xp reactivation. One hypothesis could be that PRC2-component is not recruited to the early-reactivated genes, avoiding H3K27me3 enrichment at those loci, which could enable a quick response to transcription factors such as MYC family and/or YY1 in the early ICM.

On the other hand, the presence of the repressive mark H3K27me3 on the Xp may represent a memory mark that maintains silencing at least in the later reactivated genes. In support of this hypothesis, we show that erasure of H3K27me3 during X-chromosome reactivation is at least partly an active process, as it is delayed in the absence of the H3K27 demethylase, UTX, (Figure 5). The presence of some ICM cells with complete H3K27me3 erasure in *Utx* knock-out could be explain by compensation by other demethylases such as JMJD3 and/or by passive loss of the repressive mark during cell division, however very few cell divisions occur between E3.5 and E4.5 in ICMs^44^. The interference with the kinetics of H3K27me3 loss on the Xp in *Utx* mutants correlates well with a decrease in efficiency of X-linked gene reactivation, for late reactivated genes such as *Rnf12* and *Kif4*, but not for the early-reactivated genes such as *Atrx* and *Abcb7*. This provides the first *in vivo* evidence that Utx may be involved in facilitating the Xp-reactivation process and provides important insight into the possible mechanisms involved in X-chromosome reactivation and epigenomic reprogramming in general.

In conclusion, our *in vivo* analysis of the process of Xp reactivation in the ICM reveals that different genes are reactivated by different mechanisms during ICM differentiation. Epigenetic memory of the silencing state involves H3K27me3 maintenance for some X-linked genes but not all. The reasons why some genes appear to resist full H3K27me3 during XCI and may thus be more prone to rapid reactivation, remain unknown. Interestingly, expression of several epigenetic modifiers appeared to correlate with X-chromosome reactivation (Supplementary Table 3 and Supplementary Figure 2g) such as Kdm3a, Kdm3b and Kdm3c (Jumonji C domain-containing protein that demethylates for H3K9 methylation), but also Kdm2b (H3K36-specific demethylase). MacroH2A is enriched on the inactive X chromosome^10^ and its variants (H2afy and H2afy2) are expressed in ICM cells (data not shown). MacroH2A might repress X-linked gene reactivation, in a redundant fashion with H3K27me3 marks or specifically for some genes^45^. Future work will be required to determine whether reactivation of the Xp in the ICM also requires erasure of other chromatin marks such as H3K9me2 or MacroH2A. Our findings open up the way for a better understanding of the *in vivo* requirements for epigenetic reprogramming in general.

## Methods

### Mouse crosses and collection of embryos

All experimental designs and procedures were in agreement with the guidelines from French and German legislations and institutional policies.

Mice were exposed to light daily between 7:00 AM and 7:00 PM. Noon on the day of the plug is considered as E0.5. For Figures 1 and 2, embryos were obtained by natural matings between B6D2F1 (derived from C57BL/6J and DBA2 crosses) females (5-10 weeks old) and males. For the scRNAseq experiments, hybrid embryos were derived from natural matings between C57BL/6J (B6) females (5-10 weeks old) crossed with CAST/EiJ (Cast) males.

To study the absence of Utx in early embryos, females mice carrying heterozygous or homozygous conditional *Utx* alleles (Utx^FD^, described in Thieme et al., 2014^30^) and a Cre-driven by *GDF-9* promoter (GDF9-iCre, described in Lan et al., 2004^46^) have been crossed with *Utx^FDC/Y^* males (*Utx^-/Y^)*. *Utx* control female embryos (*Utx^FDC/wt^* and *Utx^FD/FD^*) have been obtained either from the same litters as mutants (from *Utx^FD/wt^*, *GDF-9iCre* females) or after matings between *Utx^FD/FD^* females with *Utx^FD/Y^* males.

All embryos were harvested between pre-implantation to peri-implantation stages, respectively between E3.25 to E4.5. Embryos have been classified into early (E3.25-E3.5), mid (E3.75-E4.0) and late (E4.25-E4.5) blastocyst accordingly to morphology, timing and number of cells per ICM (respectively n=10-25, n=20-40 and n=30-55 cells per ICM).

### Immunosurgery for isolation of the inner cell mass

Pre-implantation blastocyst embryos at stages up to E3.5 (E4.0 for hybrid embryos) were recovered by flushing the uterus with M2 medium (Sigma). Embryos at E3.75 and later were dissected out from the uterus. The embryos were staged on the basis of their morphology and number of cells per ICM.

When applicable, the zona pellucida was removed using acid Tyrode’s solution (Sigma), and embryos were washed twice with M2 medium (Sigma). Inner Cell Mass (ICM) was then isolated from all stage blastocysts by immunosurgery as previously described^47^.

### RNA Fluorescent In Situ Hybridization

RNA FISH on blastocysts was performed as previously described^11^ using the exon and intron-spanning plasmid probe p510 for *Xist* (and its antisense *Tsix*) and BAC/Fosmid probes for genes as described in Supplementary Table 3. Images were acquired using Inverted laser scanning confocal microscope with spectral detection (LSM700 - Zeiss) equipped with a 260nm laser (RappOpto), with a 60X objective and 0.2 µm Z-sections or a 200M Axiovert fluorescence microscope (Zeiss) equippe with an ApoTome was used to generate 3D optical sections. Sequential z-axis images were collected in 0.3 µm steps. ICM obtained from *Utx^FDC/wt^* females have been PCR-genotyped after image acquisition (details available upon request).

### Immunofluorescence staining

Immunofluorescence was essentially carried out as described^48^ previously with an additional step of blocking in 3% FCS before the primary antibody incubation. All the antibodies used in this study are listed in Supplementary Table 3 along with the information on dilution ratios. Images were acquired using Inverted laser scanning confocal microscope with spectral detection (LSM700 - Zeiss) equipped with a 260nm laser (RappOpto), with a 60X objective and 0.2 µm Z-sections. Maximum projections were performed with Image J software (Fiji, NIH).

### Immunofluorescence combined with RNA Fluorescent In Situ Hybridization

Immunofluorescence followed by RNA-FISH were carried out as described previously^5^. Images were acquired using Inverted laser scanning confocal microscope with spectral detection (LSM700 - Zeiss) equipped with a 260nm laser (RappOpto), with a 60X objective and 0.2 µm Z-sections or a confocal wide-field Deltavision core microscope (Applied Precision – GE Healthcase) with a 60× objective (1,42 oil PL APO N) and 0.2 µm Z-sections or a 200M Axiovert fluorescence microscope (Zeiss) equipped with an ApoTome was used to generate 3D optical sections. Sequential z-axis images were collected in 0.3 µm steps. Images were analysed using ImageJ software (Fiji, NIH).

ICMs obtained from *Utx^FDC/wt^* females were PCR-genotyped after image acquisition (details available upon request).

All the antibodies and probes used in this study are listed in Supplementary Table 3 along with the information on dilution ratios.

### Single cell dissociation from pre-implantation to peri-implantation blastocyst stage embryos

To isolate individual cells, we incubated the ICM in TrypLE solution for 5 minutes (Invitrogen). After incubation, each blastomere was mechanically dissociated by mouth pipetting with a thin glass capillary. Single cells were then washed 3 times in PBS/acetylated BSA (Sigma) before being manually picked into PCR tubes with a minimum amount of liquid. We either directly prepared the cDNA amplification or kept the single cells at -80°C for future preparation.

### Single cell RNA amplification

PolyA^+^ mRNA extracted from each single cell was reverse transcribed from the 3’UTR and amplified following the *Tang et al* protocol^36^. Care was taken to process only embryos and single blastomeres of the highest quality based on morphology, number of cells and on amplification yield (Supplementary Table 1). Additionnal RT-specific primer for Xist amplification have been added in the lysis buffer, which contains 100nM universal RT-primer UP1 and 15nM Xist-specific RT primer ES323 (ATATGGATCCGGCGCGCCGTCGAC(T)_24_ GCAAGGAAGACAGACACACAAAGCA). Published scRNAseq samples of E3.5 trophectoderm and ICM from the same interspecific cross and the reverse cross and amplified following the same method have been added to our analysis (GSE80810; Borensztein et al., 2017^7^).

### Single cell libraries and deep-sequencing

After single cell amplification, each single cell gender has been analysed by qPCR for *Xist* and Y-linked genes *Eif2s3y*, *Uty* and *Ddx3y*. Single-cell libraries were prepared from 34 females samples, which have passed quality controls according to the manufacturer’s protocol (Illumina) and were deeply sequenced on an Illumina HiSeq 2500 instruments in single-end 50bp reads (Supplementary Table 1).

### Quality control and filtering of raw data

Quality control was applied on raw data as already described in Borensztein et al., 2017^7^. Sequencing reads characterized by at least one of the following criteria were discarded from the analysis:

1. More than 50% of low quality bases (Phred score <5).
2. More than 5% of N bases.
3. At least 80% of AT rate.
4. More than 30% (15 bases) of continuous A and/or T.

### Estimation of gene expression levels

RNA reverse transcription allowed sequencing only up to an average of 3 kb from the 3′ UTR. To estimate transcript abundance, read counts were thus normalized on the basis of the amplification size of each transcript (retrotranscribed length per million mapped reads, RPRT) rather than on the basis of the size of each gene (RPKM), as described in Borensztein et al., 2017^7^. To avoid noise due to single cell RNAseq amplification technique, only well-expressed genes (RPRT>4) were considered in our allele-specific study. A threshold of RPRT>1 was applied to consider a gene as expressed (Figures 3, 4 and Supplementary Figures 3 and 4).”

### Allele-specific RNA-seq pipeline

Allele-specific RNA-seq analysis pipeline described in Borensztein et al., 2017^7^ was applied to our data, using the same parameters, parental genomes, annotations and SNPs files. Briefly, we have filtered the SNPs on their quality values (F1 values) thanks to SNPsplit tool (v0.3.0)^49^ and SNP on chr:X 37,805,131 (mm10) in *Rhox5* gene, annotated A for C57BL/6J and G for all other strains (included C57BL/6NJ) was discarded because missing in our samples. After reconstruction of both maternal (C57BL/6J) and paternal (Castaneus) genome, allele-specific read alignment was performed with TopHat2 (v2.1.0)^50^ software. The SAMtools mpileup utility (v1.1)^51^ was then used to extract base-pair information at each genomic position. At each SNP position, the numbers of paternal and maternal alleles were counted. The threshold used to call a gene informative was five reads mapped per single SNP, with a minimum of eight reads mapped on SNPs per gene, to minimize disparity with low-polymorphic genes. The allele-specific origin of the transcripts (or allelic ratio) was measured as the total number of reads mapped on the paternal genome divided by the total number of paternal and maternal reads for each gene: allelic ratio = paternal reads/(paternal + maternal) reads.

Genes were thus classified into two categories:

1. Monoallelically expressed genes: allelic-ratio value ≤ 0.15 or ≥ 0.85.
2. Biallelically expressed genes: allelic-ratio value >0.15 or <0.85.

### Principal component analysis, hierarchical clustering and lineage analysis

Gene count tables were generated using HTSeq software (v0.6.1). Rlog function from DESeq2 R-package (v1.12.2) was used to normalize the raw counts data, with filter thresholds as described^7^. To identify the cell-origin of our samples, PCA and hierarchical clustering (Pearson correlation – Ward method) on normalised data of 23 lineage-specific factors (Figure 3) were performed using plotPCA function from DESeq2 R-package and hclust function implemented in the gplots R-package (v3.0.1) respectively.

### Heatmap of the X-chromosome

As described in Borensztein et al, 2017^7^, data from informative genes were analysed if the gene was expressed (RPRT>4) in at least 25% of the single cells (with a minimum of 2 cells except for TE) in a particular developmental stage. To follow reactivation, we decided to focus on genes at least expressed in both PrE and Epi lineages at E4.0 stage. Mean of the allelic ratio of each gene is represented for the different stages. A value has been given only if the gene was reaching the threshold described previously. Same list of genes was used for all heatmaps (116 genes). Only single cell from the same interspecific cross have been used (C57BL/6J (B6) females x CAST/EiJ (Cast) males) as different genes could follow different kinetics in a strain-specific manner^7^.

### Definition of the timing of reactivation

A minimum of 20% of expression from the Xp has been used as a threshold to call a gene as reactivated in the female samples. Adapting the method used in Borensztein et al., 2017^7^, we have automatically associated X-linked genes that become biallelic in the ICM at E3.5 (allelic ratio ≤0.15 in TE or inactivated at the same stage in Borensztein et al., 2017^7^ and >0.20 in ICM at E3.5) stage to early-reactivated gene class and in the epiblast at E4.0 stage to late-reactivated gene class (allelic ratio equals NA or ≤0.15 in TE, NA or ≤0.20 in ICM at E3.5 and >0.20 in epiblast at E4.0). X-linked genes showing very late-reactivation (0.15≤ allelic ratio in TE at E3.5 and 0.2≤ allelic ratio in other stages) in all stages are categorized as not yet reactivated genes. Finally, the last group represents genes that are escaping imprinted Xp inactivation (allelic ratio >0.15 in all stages, or NA at E3.5 and allelic ratio >0.15 in the other stages). Some genes could not be associated to a gene class due to several missing values in the decisive stages, however classes have been associated to them if RNA-FISH data was available or in case of imprinted genes (*eg Xlr3a* and *Xist* classed as “others”).

### Correlation between autosomal and X-linked gene expression

Correlation and anti-correlation between gene expression levels (autosomes and X chromosomes) and percentage of X-linked gene reactivation (allelic ratio >0.2 for X-linked genes) was measured by Pearson correlation and Benjamini-Hochberg correction and are provided in Supplementary Table 3. BC and CB (only for E3.5 trophectoderm) female single cells have been used in this analysis.

Gene ontology has been made for the top correlated genes (q-value<0.05) with the Gene Ontology Project^52^ and AMigo software^53^.

### Allele-specific H3K27me3 and H3K4me3 ChIPseq analysis

H3K27me3 and H3K4me3 enrichments in ICM were taken from Zheng et al., Mol Cell 2016^38^. Bed files of either Maternal or Paternal chromosomes for both marks were used to assess the enrichment of either marks at 5kb around their TSS. For genes having several TSS, position of start (for gene on the + strand) or end (for genes on the – strand) of the gene were taken. Score for each 100pb window containing enriched marks were sum (by Custom R scripts {R Core Team (2015). R: A language and environment for statistical computing. R Foundation for Statistical Computing, Vienna, Austria. URL https://www.R-project.org/.}). For genes whose length was below 5kb, gene size was taken as window. (Distribution of gene size for each group was not significantly different, data not shown).

### Transcription factor binding sites analysis

Nanog, Oct4, Sox2, Myc, Mycn, Klf4, Esrrb and Tcfcp2l1 binding sites from ChIPseq experiments in mouse ESCs were taken from Chen et al., Mol Cell 2008^39^. Prdm14 binding sites in mESCs were taken from Ma et al., NSMB 2011^40^. The number of binding sites of each factor in promoter, gene body and until 3kb upstream of the TSS was calculated for each gene of the reactivation-timing list (Supplementary Table 2).

### Motif discovery analysis

RSAT oligo-analysis^54^ was used to search for over-represented motifs in promoters (-700/+299nts relative to TSS) of X-linked genes in escapees, early, late and very late reactivation classes. Since the number of genes per class is too low to obtain high confidence results, we pooled genes by similar behaviour, with escapees and early reactivated genes in one group and late and very late in another one. One non-repetitive motif was found over-represented in the first group. This motif was compared to a database of known TF motifs using Tomtom (MEME Suite)^55^ and only one correspondence was found with E-value<1, that of the TF YY1 motif (p-value=0.0002, E-value=0.27, q-value=0.54). FIMO (MEME Suite)^55^ was used to determine the occurrences of this motif in each group of genes, and only matches with a p-value < 0.0001 were considered.

### Statistics section

Kruskal-Wallis and Post-hoc test were used to analyse non-parametric and unrelated samples. The statistical significance has been evaluated through two-sided Dunn’s Multiple Comparison Test with Benjamini-Hochberg correction and Kruskal-Wallis analysis of variance. p-values are provided in the figures, figure legends and/or main text. Enrichment of histone marks has been evaluated thanks to non-parametric Wilcoxon test.

### Data access

The Gene Expression Omnibus (GEO) accession numbers for the data sets reported in this paper are GSE89900 and GSE80810.

## Acknowledgements

We are grateful to P. Gestraud for help in statistical analysis. We thank the pathogen-free barrier animal facility and the Cell and Tissue Imaging Platform - PICT-IBiSA (member of France–Bioimaging) of Institut Curie. We acknowledge C.A. Penfold and the members of E.H. and A.S. laboratories for feedbacks and critical inputs. This work was funded by fellowships from Région Ile-de-France (DIM STEMPOLE), Fondation Recherche Médicale (FRM SPE20150331826) and a Marie Sklodowska-Curie Individual Fellowship (H2020-MSCA-IF-2015 - No. 706144) to M.B., CELLECTCHIP (ANR-14-CE10-0013) to E.H. and M.B, the Paris Alliance of Cancer Research Institutes (PACRI-ANR) to L.S., ERC Advanced Investigator award (ERC-2010-AdG – No. 250367), EU FP7 grants SYBOSS (EU 7th Framework G.A. no. 242129), MODHEP (EU 7th Framework G.A. no. 259743), La Ligue, Fondation de France, Labex DEEP (ANR-11-LBX-0044) part of the IDEX Idex PSL (ANR-10-IDEX-0001-02 PSL) and ABS4NGS (ANR-11-BINF-0001) to E.H, France Genomique National infrastructure (ANR-10-INBS-09) to E.H., NS, E.B., a grant-in-aid from MEXT and JST-ERATO to I.O., M.S. and a DFG grant (SPP1356) to K.A.

## Author Contributions

I.O., M.B. and E.H. conceived the study, with input from M.S., A.S and K.Anastassiadis. I.O., M.B. and K.Anastassiadis performed most of the IF/RNA-FISH experiments. K.Anastassiadis performed the IF experiments. C.P., P.D. and K. Ancelin helped for IF/RNA-FISH experiments and acquisition. M.B. performed single cell RNA amplification and C-J.C. performed the transcriptome library preparation and sequencing. L.S. and M.B. analysed the scRNAseq data and bioinformatics was supervised by NS and EB. G.G. and R.G. performed respectively the ChIPseq and the motif discovery analysis. M.B., I.O. and E.H. wrote the paper with input from all co-authors.

## Author Information

ScRNAseq data produced for this analysis are deposited in Gene Expression Omnibus under accession numbers GSE89900.

The authors declare no competing financial interests.

Correspondence and requests for material should be addressed to E.H.

## Supplemental Information

**Supplementary Figure 1 (related to Figure 4).**
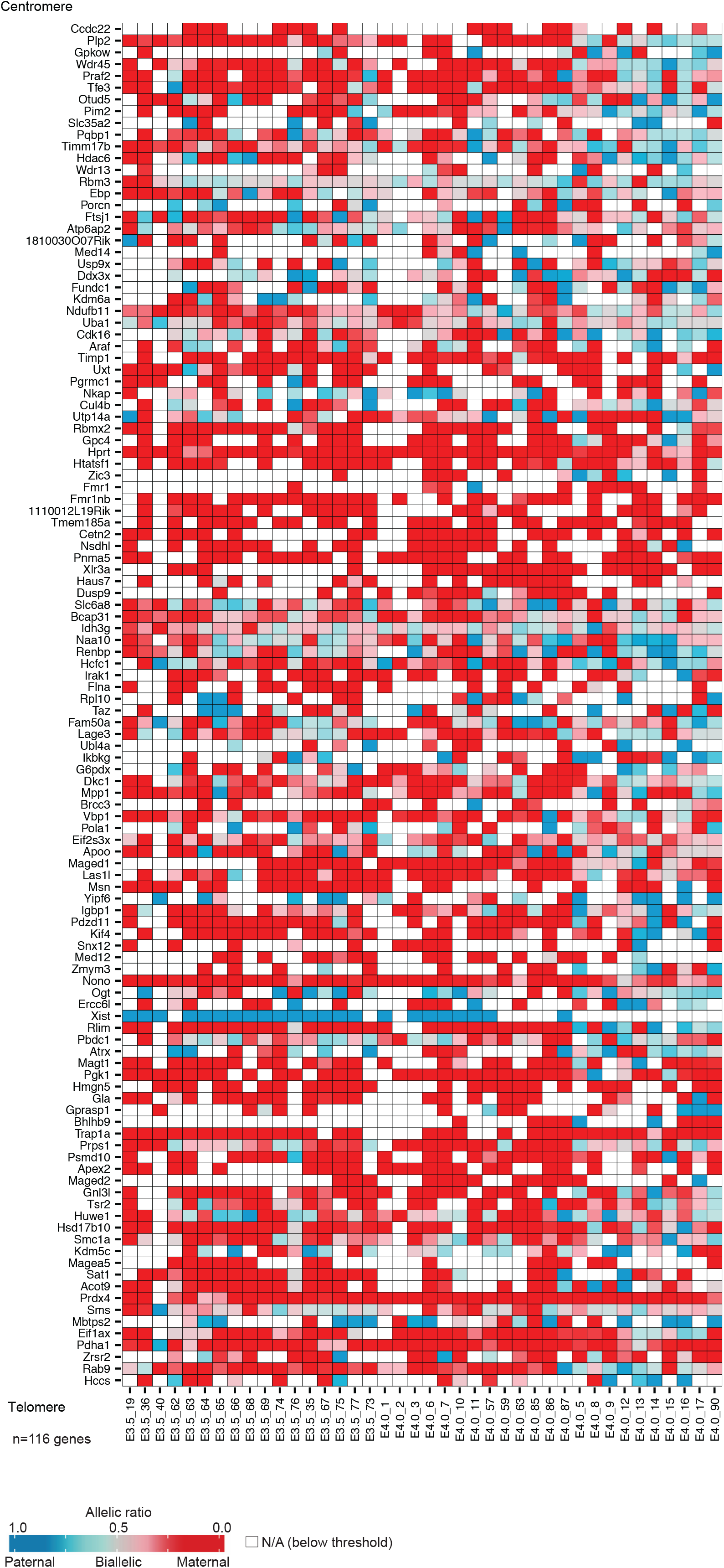
Heatmap representing the allele-specific expression of informative and well expressed X-linked genes in each single cell, in E3.5 (Trophectoderm and ICM) and E4.0 (Primitive Endoderm and Epiblast) female hybrid embryos (B6 x Castaneus). Strictly maternally expressed genes (allelic ratio ≤0.15) are represented in red and strictly paternally expressed genes (allelic ratio ≥0.85) in blue. Colour gradients are used in between and genes have been ordered by genomic position. n=116 genes.

**Supplementary Figure 2 (related to Figure 4).**
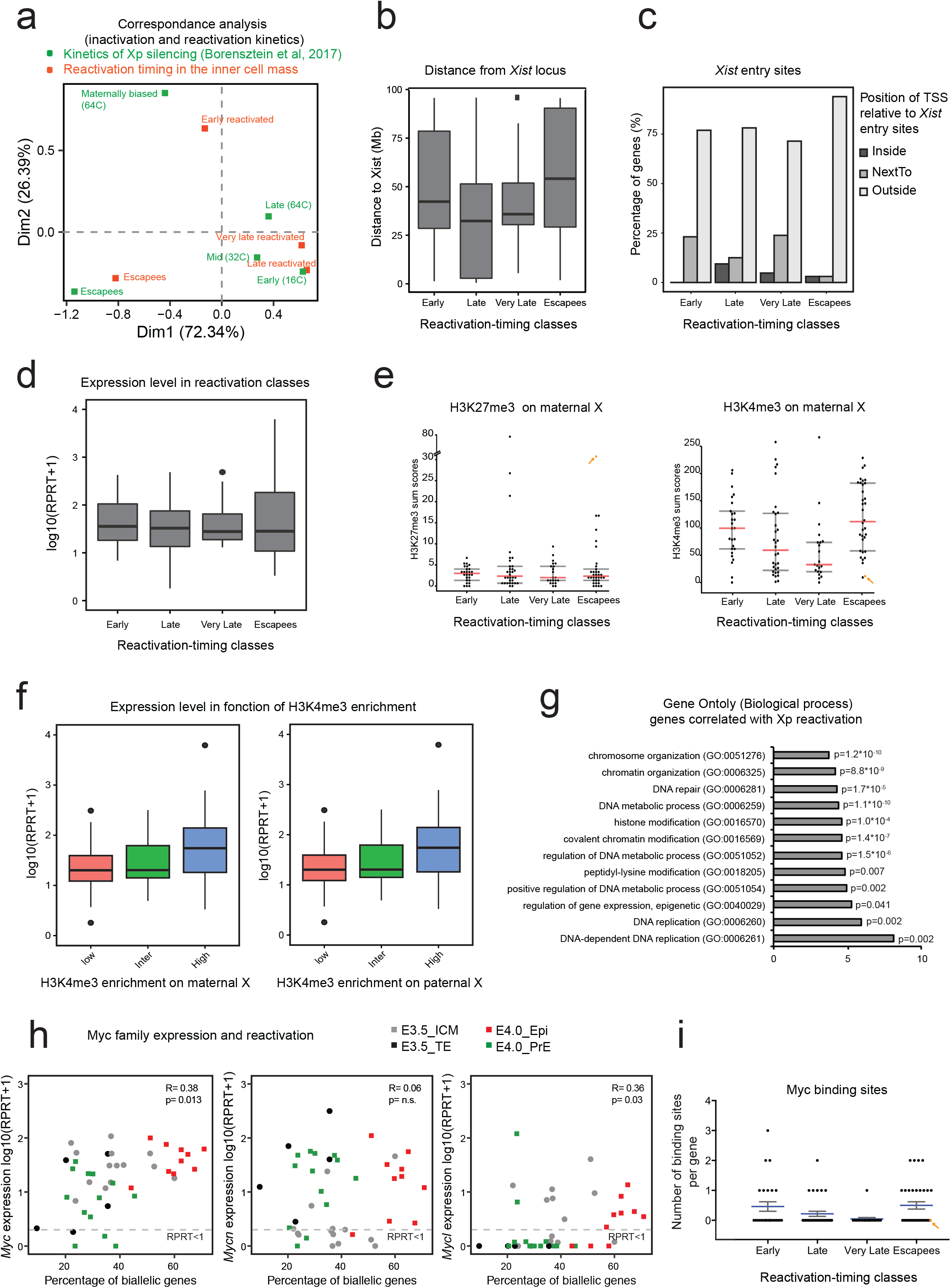
**(a)** Correspondence analysis (CA) of X-linked gene reactivation and silencing classes based on their timing of reactivation in ICM and timing of silencing during imprinted XCI in pre-implantation embryos as previously determined in Borensztein *et al*, 2017^7^. **(b)** Distance to *Xist* genomic locus. Distribution of the genomic distances to *Xist* locus (in Mb) for the different X-linked gene reactivation classes. Transcription Start Site (TSS) of each gene has been used to measure the distance to *Xist* locus. Non-significant by Kruskal-Wallis test. Boxplot represent median with lower and upper quartiles. **(c)** Percentage of X-linked genes from the different reactivation classes classified by their relative position to *Xist* “entry” sites (as identified during XCI induction in ESCs^37^: “inside” (TSS located in a *Xist* “entry” site), “next to” (TSS located less than 100 kb to an “entry” site) and “outside” (over 100 kb from an “entry” site). Non-significant by Kruskal-Wallis test. **(d)** Expression level of X-linked genes in the different reactivation-timing classes in E3.5 ICM samples (mean of each single gene). No differences in expression level can be seen for early reactivated and escapees genes compared to late and very late genes. Boxplot represent median with lower and upper quartiles. Non-significant by Kruskal-Wallis test. **(e)** Enrichment of H3K27me3 and H4K4me3 on maternal X obtained from (Zheng et al., 2016)^38^. Each dot represents a single gene. *Xist* dot is highlighted with an orange arrow. No differences can be seen for H3K27me3 distribution in any reactivation-timing groups (by Wilcoxon test), contrary to the paternal X (Figure 4c). Enrichment of H3K4me3 is much higher on maternal X chromosome compared to all paternal X (Figure 4c). Very late genes are significantly different compared to Early and Escapee groups for H3K4me3 maternal enrichment by Wilcoxon test (respectively p=3.61*10^-3^ and p=1.71*10^-4^). **(f)** Expression level of X-linked genes in function of their enrichment of H3K4me3 on maternal (left) and paternal (right) X chromosomes. Low, intermediate (Inter) and highly (High) enriched classes have been designed by H3K4me3 sum scores <5, 5≤ and ≥ 15, and >15 respectively. On the maternal and paternal X chromosomes, lowly enriched genes for H3K4me3 marks are significantly less expressed than highly expressed ones (respectively p=0.028 and p=0.045, by Dunn’s test). Boxplot represent median with lower and upper quartiles. **(g)**Representation of the Gene ontology analysis of Biological process performed on the best correlated genes with X-linked gene reactivation (q-value <0.05, Supplemental Table 3). The twelve best enrichment classes (based on fold enrichment) are represented with their p-value. **(h)**The level of expression of *Myc* genes (*Myc, Mycn* and *Mycl*) is plotted in function of the number of biallelically/reactivated X-linked genes in each single cell. Different colours are applied for E3.5 trophectoderm (TE), E3.5 ICM, E4.0 Primitive endoderm (PrE) and E4.0 Epiblast (Epi) cells. By Spearman correlation, a positive correlation is seen between level of expression of *Myc* and *Mycl* and high percentage of biallelically expressed genes. Genes with level of expression as (RPRT<1) are considered as non-expressed in our samples. **(i)**Mean (+/-s.e.m.) of the number of Myc family (Mycn and Myc) binding sites per gene in each reactivation-timing groups, obtained from Chen et al., 2008^39^. There is a significantly higher number of genes with at least one binding site for Myc factors in early and escapee groups, p=0.0269 by Kruskal-Wallis test. *Xist* is highlighted with an orange arrow.

**Supplementary Figure 3 (related to Figure 5).**
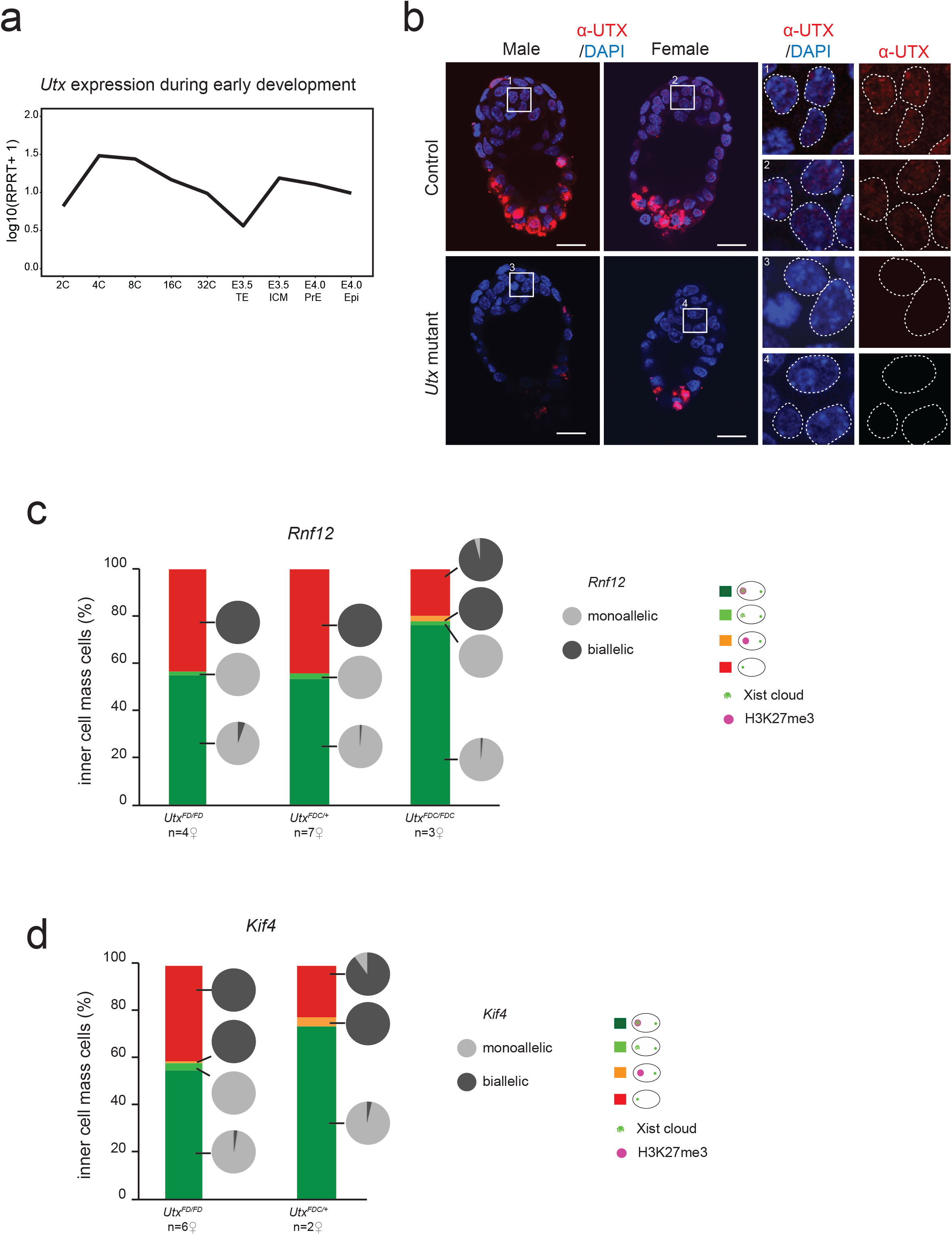
**(a)** Level of expression of *Utx* gene during preimplantation mouse development (expression mean of all single cells). *Utx* is downregulated in trophectoderm but stay expressed in ICM cells at E3.5. **(b)** Maximum intensity projection of 1.5 µm section for control *(Utx^FD/FD^* female and *Utx^FD/Y^* male) and mutant (*Utx^FDC/FDC^* female and *Utx^FDC/Y^* male) E4.5 blastocysts analysed by immunofluorescence against UTX (red). DAPI is in dark blue. Enlarged nuclei are shown. Scale bars represent 20µm. **(c)**Proportion (mean) of ICM cells showing enrichment of H3K27me3 on the Xist RNA coated X chromosome from E4.5 control *(Utx^FD/FD^*), heterozygous *(Utx^FDC/+^)* and mutant (*Utx^FDC/FDC^*) female blastocysts, linked with *Rnf12* allelic status. **(d)**Proportion (mean) of ICM cells showing enrichment of H3K27me3 on the Xist RNA coated X chromosome) from E4.5 control *(Utx^FD/FD^*) and mutant (*Utx^FDC/FDC^*) female blastocysts, linked with *Kif4* allelic status.

**Supplementary Table 1: Summary of single cell RNAseq samples.** For each library is provided: single cell’s name, stage, embryo number, gender, cross and the raw read number, filtered ones and percentage of mapping.

**Supplementary Table 2: Silencing gene classes** Reactivation timing and allelic ratio for the 116 informative and well-expressed X-linked genes in hybrid ICM cells (B6 x Cast cross).

**Supplementary Table 3: List of genes correlated or anti-correlated with X-linked gene reactivation between E3.5 and E4.0.**

**Supplementary Table 4: List of RNA-FISH probes and antibodies**

